# Competition for iron shapes metabolic antagonism between *Bacillus subtilis* and *Pseudomonas*

**DOI:** 10.1101/2023.06.12.544649

**Authors:** Mark Lyng, Johan P. B. Jørgensen, Morten D. Schostag, Scott A. Jarmusch, Diana K. C. Aguilar, Carlos N. Lozano-Andrade, Ákos T. Kovács

## Abstract

Siderophores have long been implicated in sociomicrobiology as determinants of bacterial interrelations. For plant-associated genera like *Bacillus* and *Pseudomonas*, siderophores are well known for their biocontrol functions. Here, we explored the functional role of the *Bacillus subtilis* siderophore bacillibactin in an antagonistic interaction with *Pseudomonas marginalis*. The presence of bacillibactin strongly influenced the outcome of the interaction in an iron-dependent manner. The bacillibactin producer *B. subtilis* restricts colony spreading of *P. marginalis* by repressing the transcription of histidine kinase-encoding gene *gacS*, thereby abolishing production of secondary metabolites such as pyoverdine and viscosin. By contrast, lack of bacillibactin restricted *B. subtilis* colony growth in a mechanism reminiscent of a siderophore tug-of-war for iron. Our analysis revealed that the *Bacillus-Pseudomonas* interaction is conserved across fluorescent *Pseudomonas* spp., expanding our understanding of the interplay between two genera of the most well-studied soil microbes.

## INTRODUCTION

Interactions between bacteria is of huge relevance to how microbial ecology can be manipulated for biotechnological benefit. In natural environments, bioavailable iron is often scarce and because it is an essential mineral to virtually all life, many organisms have evolved to scavenge iron using high-affinity metal chelators [1]. Metal chelators are a main driver of antagonism between microorganisms both in sequestering metal ions from competing organisms, and through bioactive toxicity directly inhibiting or even killing competitors [2, 3].

The catecholate siderophore bacillibactin (BB) produced by members of the *Bacillus* genus is an example of an iron chelator that performs multiple functions. In *Bacillus subtilis*, it is the main iron acquisition metabolite with high affinity for ferric iron, Fe(III) [4]. BB is synthesised through a non-ribosomal peptide synthetase pathway encoded by the *dhbA−F* operon and is exported to acquire Fe(III) before being taken up again by the FeuABC-YusV ABC transporter system [5–8]. Once intracellular, BB-Fe is hydrolysed by BesA to release iron ions for use in enzymatic reactions and in transcriptional regulation through the ferric uptake regulator Fur [7].

However, the *dhbA−F* operon also plays an essential role in *Bacillus* biofilm formation. The *ΔdhbA* mutant lacks dehydrogenation of (2S,3S)-2,3-dihydroxy-2,3-dihydrobenzoate to 2,3-dihydroxybenzoate (DHB) and therefore is unable to produce pellicle biofilms in biofilm-promoting defined medium MSgg due to an inability to perform extracellular electron transport with ferric iron captured by DHB [9]. Additionally, BB from *Bacillus amyloliquefaciens* has even been suggested to exhibit direct bioactivity against the plant-pathogenic *Pseudomonas syringae* pv. *tomato*, though without direct experimental evidence [10], and recent studies have demonstrated antibacterial properties of BB isomers [11, 12].

*Bacillus* and *Pseudomonas* are bacterial genera with high environmental impacts. They are environmentally ubiquitous and often co-isolated, especially within the context of crops, where they function as biostimulants and biocontrol agents, but also, in the case of *Pseudomonas*, as phytopathogens [13]. Both genera contain members capable of producing bioactive natural products that antagonise competing microbes [14–16], and both can alter the local microbiome [17, 18]. Such abilities have immense implications in areas such as agriculture, where microbiome composition is consistently correlated with plant health and disease outcome [19, 20], and therefore considered an environmentally friendly agricultural practice.

Pairwise interactions between bacilli and pseudomonads might therefore play a crucial role in determining plant growth, health, and local microbiome composition. Studies investigating such pairwise interactions are revealing more detail, outlining the molecular mechanisms, and determining the degree to which interactions and their effector molecules are conserved [21–25]. However, despite these advances, the large bioactive potential of *Bacillus* and *Pseudomonas* makes mapping the overarching interaction mechanisms challenging, and a consensus has not been reached on the general interactions between members of these two genera. Some studies investigated the role of iron and siderophores in interactions between them, identifying *Pseudomonas* siderophores pyoverdine (Pvd) and pyochelin as signalling molecules for *Bacillus velezensis* [21] and pulcherriminic acid as an iron sequestering agent in interactions between *B. subtilis* and *Pseudomonas protegens* [26].

Here, we investigated the functional role of *B. subtilis* BB in antagonising *Pseudomonas*. Using a macrocolony competition assay, we observed that *B. subtilis* DK1042 (hereafter DK1042) was unable to restrict the colony size of *Pseudomonas marginalis* PS92 (hereafter PS92) when lacking the complete BB biosynthetic pathway, and that products of DhbA−F enzymes can be secreted and shared among *B. subtilis* cells. The interaction depended on iron in the media, but transcriptomic analysis revealed no iron starvation response in cocultures. Instead, the PS92 Gac/Rsm signalling system, herein Pvd biosynthesis, was strongly downregulated transcriptionally and metabolically when cocultured alongside DK1042. Finally, the specific interaction of DK1042 and the consequence of lacking BB were found to be conserved among fluorescent pseudomonads.

## METHODS

### Culturing

*B. subtilis* DK1042 (the naturally competent derivative of 3610) and *Pseudomonas marginalis* PS92 were routinely cultured in lysogeny broth (LB; Lennox, Carl Roth, 10 g/L tryptone, 5 g/L yeast extract and 5 g/L NaCl), LB supplemented with 1% (v/v) glycerol and 0.1 mM MnCl_2_ (LBGM), or King’s B medium (KB; 20 g/L peptone, 1% glycerol (v/v), 8.1 mM K_2_HPO_4_, 6.08 mM MgSO_4_·7H_2_O) at 30°C. Colonies were grown on media solidified with 1.5% agar (w/v). FeCl_3_, ammonium ferric citrate and 2,2’-bipyridine (BPD) were added to autoclaved agar at varying concentrations. Antibiotics were added in the following final concentrations: 50 μg/mL gentamycin (Gm), 100 μg/mL ampicillin (Amp), 5 μg/mL chloramphenicol (Cm), 10 μg/mL Kanamycin (Km).

### Pairwise interactions on agar

DK1042 and PS92 were routinely spotted 5 mm apart on agar surfaces using 2 µL culture at an optical density at 600 nm (OD_600_) of 1.0. Prior to spotting, plates were dried for 30 min in a lateral flow hood then incubated at 30°C. When spotting cocultures, each species was adjusted to an OD_600_ of 1.0 and the two strains were then mixed in equal volumes. For mixed *B. subtilis* cultures, wild-type (WT) and mutant strains were mixed 1:1, 1:10, or 1:100, based on OD_600_.

### Stereomicroscopy

Interactions for stereomicroscopy were assessed using DK1042 carrying *amyE::*P*_hyperspank_*-*mKate2* and PS92 carrying *attTn7::msfGFP*. Colonies were imaged with a Carl Zeiss Axio Zoom V16 stereomicroscope equipped with a Zeiss CL 9000 LED light source and an AxioCam 503 monochromatic camera (Carl Zeiss, Jena, Germany). The stereoscope was equipped with a PlanApo Z 0.5×/0.125 FWD 114 mm, and filter sets 38 HE eGFP (ex: 470/40, em: 525/50) and 63 HE mRFP (ex: 572/25, em: 629/62). Exposure time was optimised for appropriate contrast.

Image processing and analysis were performed using FIJI (2.1.0/1.53f51) [27]. Contrast in fluorescence channels was adjusted identically on a linear scale. Colony area was measured by segmenting the colony of interest using Otsu’s algorithm [28] based on fluorescence (msfGFP or mKate2 for *Pseudomonas* or *Bacillus*, respectively).

### Genetic modification

DK1042 mutants were created through homologous recombination using the *B. subtilis* single gene deletion library [29]. In brief, genomic DNA from a donor *B. subtilis* 168 mutant was extracted using a EURx Bacterial & Yeast Genomic DNA Purification Kit (EURex, Gdansk, Poland) following the manufacturer’s instructions, and 100−200 ng genomic DNA was added to DK1042 grown to OD_600_ ∼0.5 in 400 µL at 37°C in competence medium (80 mM K_2_HPO_4_, 38.2 mM KH_2_PO_4_, 20 g/L glucose, 3 mM triNa citrate, 45 µM ferric NH_4_ citrate, 1 g/L casein hydrolysate, 2 g/L K-glutamate, 0.335 µM MgSO_s_·7H_2_O, 0.005 % [w/v] tryptophan). Cells were incubated with DNA at 37°C for 3 h before 100 µL of the transformation mix was spread onto LB agar supplemented with kanamycin and incubated at 37°C for 16−24 h. Mutants were validated by colony PCR with primers designed to anneal 50−150 bp up- and downstream of the gene of interest (Table S1).

PS92 attTn7::*msfGFP* was constructed by conjugation with pBG42 as described by Zobel *et al.* (2015) [30]. In brief, cultures of *Escherichia coli λpir* CC118/pBG42 (donor - Gm^R^), *E. coli* HB101/pRK600 (helper, Cm^R^), *E. coli λpir* CC118/pTNS2 (transposase-carrying, Amp^R^) and PS92 (recipient) were started in LB medium supplemented with appropriate antibiotics and incubated overnight. Each culture was washed thrice in 0.9% NaCl before being mixed equally based on OD_600_. The mix was spotted onto LB without antibiotics and incubated overnight at 30°C. Resultant colonies were resuspended in 0.9% NaCl and plated on Pseudomonas Isolation Agar (45.03 g/L; Millipore, Burlington, Massachusetts, United States) supplemented with Gm to select for positive *Pseudomonas* conjugants.

### Mass spectrometry imaging

Pairwise interaction colonies were grown on 10 mL agar plates incubated for 72 h before being excised, dried, and sprayed for matrix-assisted laser desorption ionization-mass spectrometry imaging (MALDI-MSI). Agar was adhered to MALDI IntelliSlides (Bruker, Billerica, Massachusetts, USA) using a 2-Way Glue Pen (Kuretake Co., Ltd., Nara-Shi, Japan). Slides were covered by spraying 1.5 mL of 2,5-dihydrobenzoic acid (40 mg/mL in ACN/MeOH/H_2_O [70:25:5, v/v/v]) in a nitrogen atmosphere and dried overnight in a desiccator prior to MSI acquisition. Samples were then subjected to timsTOF flex using a Bruker Daltonik GmbH mass spectrometer for MALDI MSI acquisition in positive MS scan mode with a 40 µm raster width and a mass range of 100−2000 Da. Calibration was performed using red phosphorus. The settings in timsControl were as follows: Laser: imaging 40 µm, Power Boost 3.0%, scan range 46 µm in the XY interval, laser power 70%; Tune: Funnel 1 RF 300 Vpp, Funnel 2 RF 300 Vpp, Multipole RF 300 Vpp, isCID 0 eV, Deflection Delta 70 V, MALDI plate offset 100 V, quadrupole ion energy 5 eV, quadrupole loss mass 100 m/z, collision energy 10 eV, focus pre TOF transfer time 75 µs, pre-pulse storage 8 µs. After data acquisition, data were analysed using SCiLS software (version 2021b Pro).

Images were prepared from data normalised to the root-mean square. Images from each m/z value were expanded to be of equal size by adding pixels with NaN values if necessary. Linear scaling of pixel intensity was performed across images displaying the same m/z values. NaN is presented as 0 in the final images.

### Liquid chromatography mass spectrometry (LC-MS)

*B. subtilis* DK1042 was inoculated into 20 mL LB in a 100 mL flask and grown overnight. The culture was adjusted to OD_600_ = 1.0 and 1 mL was spread on each of 150 LBGM agar plates and incubated at 30°C for 72 h. Extraction and subsequent LC-MS analysis were performed as previously described [31].

### Whole-genome sequencing and genome comparisons

*Pseudomonas* genomes were sequenced and assembled using a hybrid approach with Oxford Nanopore and Illumina sequencing. From a culture grown in LB overnight, genomic DNA was extracted using an EURx Bacterial & Yeast Genomic DNA Purification Kit following the manufacturer’s instructions. Short read library preparation and sequencing were performed at Novogene Co., Ltd. on an Illumina NovaSeq PE150 platform. Long-read library preparation was performed using an SQK-RBK004 Oxford Nanopore Rapid Barcoding kit, libraries were sequenced on a fresh Oxford Nanopore MinION Mk1B 9.4 flow cell for 12 h, and base called simultaneously with Guppy in MinKNOW (Oxford Nanopore Technologies, Oxford United Kingdom) using the default r941_min_hac_g507 model. Genomes were assembled using Trycycler (V0.5.3) [32] when the long read sequencing depth was adequate for subsetting or Unicycler (V0.5.0) [33] in all other cases. Adapters were removed using Porechop (0.2.4 https://github.com/rrwick/Porechop) and reads were filtered with Filtlong (V0.2.1 https://github.com/rrwick/Filtlong). Default Trycycler and Unicycler workflows were applied to assemble genomes (see Supplementary Methods for details). Completeness and contamination were assessed using CheckM (V1.1.3) [34] and whole-genome taxonomy was determined with AutoMLST [35] and the TYGS database (V342) [36].

Assembled genomes were annotated with Bakta (V1.6.1) [37] and a gene presence/absence matrix was created with Panaroo (V1.3.2) [38] using its moderate mode and a CD-HIT amino acid identity threshold at 40%.

### Transcriptomics

Colonies were collected with a 10 µL loop and placed in RNAse-free screw-cap tubes to sample either the interaction zone (∼2 mm into each colony for pairwise interactions) or the entire colony (for mono-cultures). The collected biomass was solubilised in RNAProtect and frozen in liquid nitrogen before being stored at −80°C prior to RNA purification, for no more than 3 days.

RNA was extracted using a Qiagen RNeasy PowerMicrobiome RNA extraction kit (QIAGEN N.V., Venlo, The Netherlands) via combined bead beating and phenol-chloroform extraction (see Supplementary Methods for details). Ribosomal RNA depletion, library preparation and sequencing were performed at Novogene Co., Ltd. on an Illumina NovaSeq PE150 platform.

Read abundances were determined using Kallisto (0.48.0) [39]. The genome of PS92 was annotated with Bakta (V1.6.1) [37] while the genome of *B. subtilis* DK1042 (Acc: CP020102 and CP020103) was retrieved from GenBank. Genomes were converted into a fasta-formatted transcript file with gffread (v0.11.7) [40] then combined into a Kallisto index.

RNA sequencing (RNA-seq) datasets were trimmed and filtered with FastP, and reads were quantified by mapping every sample to the combined index file (consisting of transcripts from both species), with 100 bootstrap replicates. Differential expression analysis was performed with DESeq2 (v1.34.0) [41] in Rstudio (2022.02.3-b492) [42] running R (4.1.1) [43] with the Tidyverse framework (1.3.1) [44]. Pathway analysis was performed by functionally annotating the transcriptome of each species using eggNOG-mapper V2 (online browser version) with default parameters [45]. Differentially expressed pathways were investigated using KEGG Mapper Reconstruct [46], and manually using the databases Subtiwiki [47] and The *Pseudomonas* Genome Database [48] as references.

## Data availability

Analysis scripts and processed data have been deposited at Github (https://github.com/mark-lyng/dhb_story).

Genome assemblies have been deposited at NCBI (accession numbers are listed in Table S1).

Raw sequence reads (long- and short-gDNA reads and RNA) have been deposited at the Sequencing Read Archive (SRA; https://www.ncbi.nlm.nih.gov/sra) with BioProject ID PRJNA956831.

Raw microscopy images have been deposited at Dryad (https://datadryad.org/; DOI https://doi.org/10.5061/dryad.vq83bk3zc).

MSI data have been deposited at Metaspace [49] under project ID https://metaspace2020.eu/pro-ject/lyng-2023

LC-MS data have been deposited at GNPS-MassIVE under MSV000092142.

## RESULTS

### BB is involved in mutual antagonism between DK1042 and PS92

Molina-Santiago and colleagues previously described that on solid KB medium (a growth medium for *Pseudomonas*-produced Pvd detection), DK1042 surrounds and restricts the growth of *Pseudomonas chlororaphis* PCL1606, unlike on various other media used in their experiments [24]. Therefore, we tested a soil-isolated *Pseudomonas* species (designated PS92) and revealed an interaction phenotype comparable to that of PCL1606. Since antagonism was apparent on KB medium, which induces Pvd production in *Pseudomonas*, and not on LB medium (Fig. S1a), we hypothesised that BB, the catechol siderophore of *B. subtilis*, is involved in this interaction. To evaluate the effect of BB in the interaction between DK1042 and PS92, we cultured fluorescently labelled versions of the two species together in close-proximity macro colonies on solid agar medium. Spots (2 µL) of each culture were placed 5 mm apart and grown at 30°C for 3 days (Fig. 1a and Video S1). We compared WT DK1042 and the single gene disruption mutant (Δ*dhbA*) lacking the ability to produce DHB and BB.

**Fig 1.**
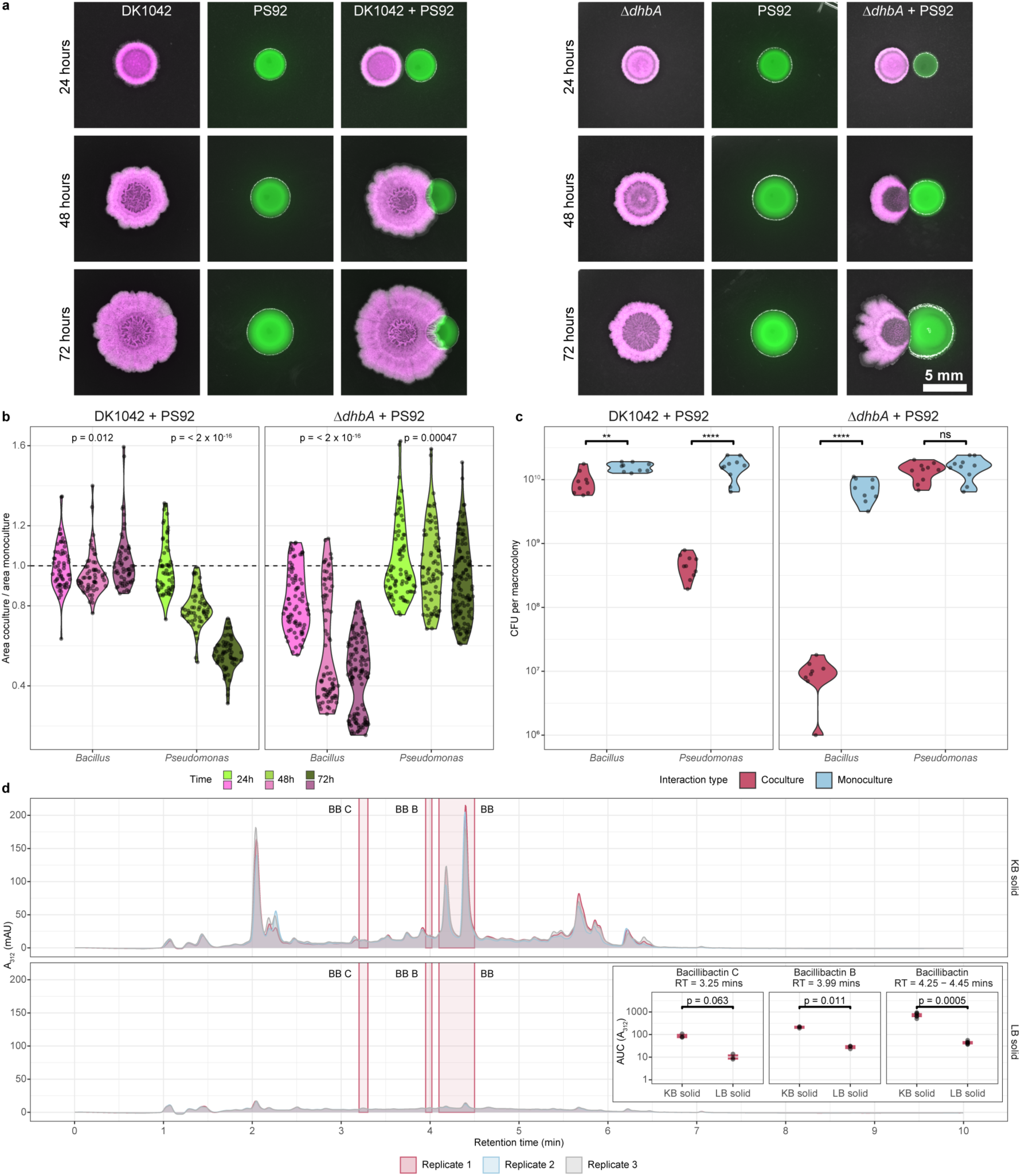
Antagonism is DhbA-dependent. **a:** DK1042 (mKate2, magenta) and PS92 (msfGFP, green) were cocultured on KB agar with 5 mm between colony centres and imaged every 24 h. Scale = 5 mm. **b:** Coculture colony area was normalised against monoculture colony area and plotted for each partner (x-axis) in the two interactions (facets). The dashed line indicates coculture = monoculture. The *p*-value was calculated by analysis of variance (ANOVA) within groups (n = 40 colonies from two independent experiments). **c:** Colony-forming units were determined from the macrocolonies in coculture and monoculture. Significance stars are from Student’s t-tests (n = 10 colonies from two independent experiments; \*\**p* <0.01, \*\*\*\**p* <0.0001; ns: not significant. **d:** Ultra-violet quantification of LC-MS on DK1042 monocultures at λ = 312 nm. Bacillibactin and isomers in KB (above) and LB (below). BB, Bacillibactin; BB B, Bacillibactin B; BB C, Bacillibactin C. Inset: Peak area under the curve (AUC) of bacillibactin and isomers in KB and LB. Labels are *p*-values from Student’s t-test (n = 3 independent experiments).

By measuring the areas of colonies, we determined that the growth of PS92 cocultured next to DK1042 was restricted and after 72 h; the area of monoculture colonies was approximately twice that of the area of coculture colonies (Fig. 1b). The Δ*dhbA* mutant did not surround nor restrict the growth of PS92, but rather, was itself restricted approximately two-fold. The restriction in area is reflected in the number of colony-forming units from colonies in each interaction, showing a 16-fold reduction in cells in PS92 colonies grown next to DK1042 and a 10^3^-fold reduction in cells from Δ*dhbA*-mutant colonies grown next to PS92 (Fig. 1c). When coculturing the strains in liquid medium, we observed inhibition of PS92 in KB broth regardless of whether the coculture was with DK1042 or Δ*dhbA*. Neither DK1042 nor Δ*dhbA* was antagonised by PS92, demonstrating that the observed antagonism is specific to solid media (Fig. S1d and S1e). We also observed that the interaction occurs on LBGM but not on MSgg, and that glycerol is likely implicated in the antagonism (Fig. S2).

In previous studies, BB production was reported to be linked to iron stress conditions [10]. As both LB and KB contain sources of natural iron from tryptone or yeast extract, low BB production could be expected on both types of media. LC-MS was used to compare BB production from DK1042 inoculated in liquid and solid LB and KB, revealing a 10-fold increase in BB concentration on solid KB compared to solid LB (Fig. 1d), while virtually undetectable in liquid media (Fig. S3). The dependency on *dhbA* for antagonism and defence coupled with the exclusive production on solid KB suggests a BB-mediated regulation in DK1042. However, a previous study showed that in *B. subtilis* biofilms, BB but not DHB is dispensable in air-liquid interface pellicle biofilms [9]. Thus, we tested whether BB or any precursor from the *dhbA-F* pathway influenced the pairwise interaction (Fig. 2a) and found all mutants unable to antagonise PS92 (Fig. 2b), suggesting that BB, and not an intermediate, must be present for antagonism.

**Fig 2.**
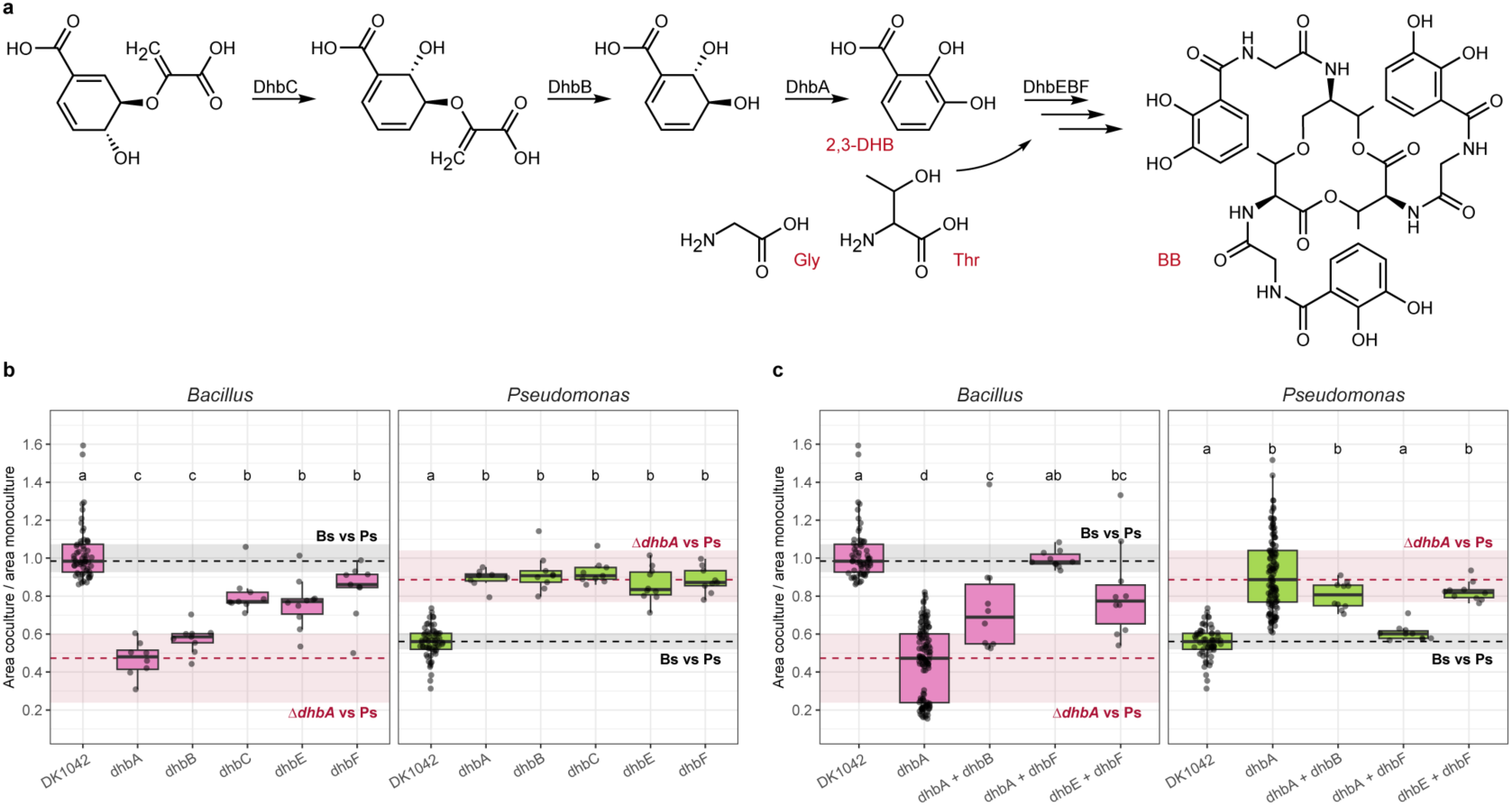
Bacillibactin precursors are ‘common goods’. **a:** Bacillibactin (BB) is produced by the bio-synthetic enzymes DhbACEBF from chorismate to 2,3-dihydroxybenzoate (2,3-DHB), which is then used as a starting unit by the non-ribosomal synthase complex DhbEBF to cyclize three monomers of 2,3-DHB using glycine (Gly) and threonine (Thr) linkers. **b:** Relative colony areas of PS92 (green) and individual Δ*dhbA−F* gene deletion mutants (magenta). **c:** Relative colony areas of PS92 (green) and combinations of Δ*dhbA−F* gene deletion mutants (magenta). Black dashed lines and grey rectangles denote the mean relative colony size in the DK1042 + PS92 interaction (lines = mean, rectangles = min and max). Red dashed lines and rectangles denote similarly for the Δ*dhbA* + PS92 interaction. Grouping letters are from ANOVA with Tukey-Kramer’s Post-hoc test. Identical letters within each plot indicate a statistically significant grouping (*p* <0.05, n >8, from three independent experiments).

### Bacillibactin precursors are ‘common goods’

Since DHB has been shown to function as a siderophore in biofilms [50], we hypothesised that it could be used as a ‘common good’ and taken up by Δ*dhbA* mutant cells. Therefore, we mixed mutants disrupted in early biosynthesis (Δ*dhbA* and Δ*dhbB*), late biosynthesis (*ΔdhbE* and *ΔdhbF*), and mutants disrupted in both early and late biosynthesis (Δ*dhbA* and Δ*dhbF*) that should be able to complement each other through Δ*dhbF* supplying DHB to Δ*dhbA*, which then completes the synthesis of BB. Only the mixture of Δ*dhbA* and Δ*dhbF* was able to antagonise and restrict the growth of PS92 on solid KB medium (Fig. 2c), which not only demonstrates the ability of the mutants to complement each other, but also confirms that BB, and not DHB, is responsible for the antagonistic interaction between DK1042 and PS92. This was further corroborated by mixing DK1042 with Δ*dhbA* (unable to produce DHB) or Δ*dhbF* (unable to synthesise the non-ribosomal peptide) using 1:1, 1:10 and 1:100 ratios, respectively, and spotted mixtures next to PS92 (Fig. S4). Both mutants were rescued by the WT strain, even at a ratio of 1:100, and expansion of PS92 colonies was restricted independently of which mutant was used in the mixture or the ratio of the mixed strains. This indicates that the critical components of the DhbA-F pathway can be secreted and shared between members in a population.

### Excess iron abolishes antagonism

Given that BB is the main siderophore of *B. subtilis*, we hypothesised that the antagonism was driven by iron in the media. To test this hypothesis, we cocultured DK1042 and PS92 on KB agar supplemented with varying concentrations of the ferrous iron chelator BPD (Fig. 3a) or with excess ferric iron either as ferric citrate, which is not transported into the cell via the BB and FeuABC-YusV machinery (Fig. 3b), or as FeCl_3_ from which iron is readily bound to DHB and BB (Fig. 3c) [6, 9]. Curiously, sequestration of Fe(II) and addition of Fe(III) resulted in increased coculture colony size for the Δ*dhbA* mutant. Sequestering ferrous iron did not lead Δ*dhbA* to regain the ability to restrict PS92 size, but, independently of concentration, both the addition of FeCl_3_ and ferric citrate did, even though Δ*dhbA* should be unable to utilise FeCl_3_ [51]. Similarly, the Δ*besA* mutant unable to release iron from BB was also complemented by any iron-related change to the medium. However, the Δ*feuA* mutant did not respond to FeCl_3_ (as expected), but could be complemented by ferric citrate in a dose-dependent manner. Though the iron concentration added to the medium in these experiments was relatively high, these results suggest that the interaction, both defensive and offensive, is iron-dependent.

**Fig 3.**
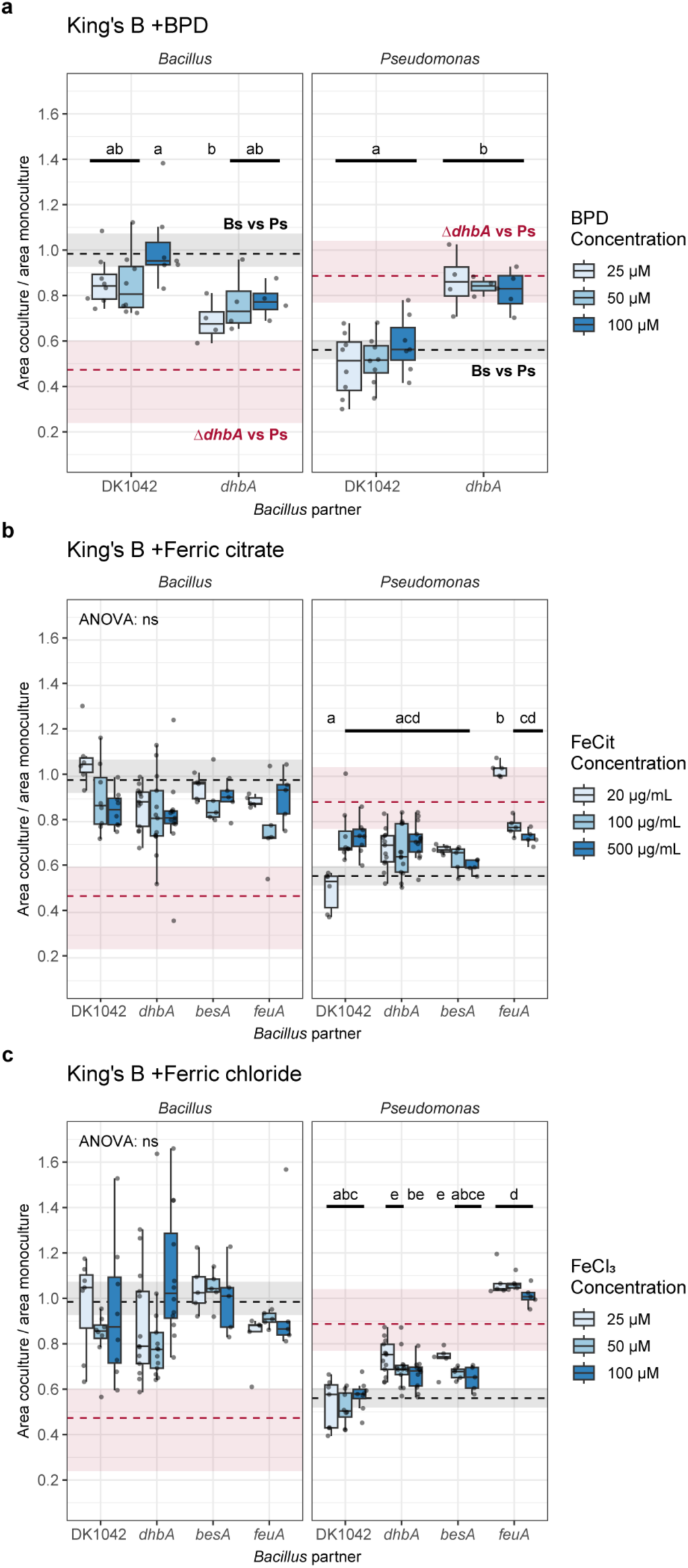
Excess iron restores mutant bacilli offensive and defensive attributes. *Bacillus* and *Pseudomonas* relative colony areas on KB with increasing concentrations of **a:** Fe(II) chelator 2,2-bipryidine (BPD), **b:** ferric citrate (FeCit) and **c:** ferric chloride (FeCl_3_). PS92 was spotted next to DK1042 or mutants deficient in the bacillibactin biosynthesis and uptake pathway. Dashed lines and rectangles are median, 1st and 3rd quartiles for PS92 colonies spotted next to Δ*dhbA* (red) and DK1042 (grey) on non-supplemented KB. Grouping letters are from ANOVA with Tukey-Kramer’s Post-hoc test. Identical letters within each plot indicate a statistically significant grouping (*p* <0.05, n ≥4 colonies from two independent experiments).

### DK1042 heavily alters the PS92 transcriptome

To better understand the transcriptional landscape present in each strain as a result of cocultivation, we extracted total RNA from colonies cultured in monoculture or with an interaction partner. This allowed us to identify transcriptional differences between DK1042 and Δ*dhbA* both in monoculture and coculture, but also between each individual strain in monoculture compared to coculture to ascertain how a neighbouring partner alters the transcription of a focal strain (Fig. S5). Transcriptomic analysis revealed that the PS92 transcriptome was heavily altered by DK1042 (1036 differentially regulated genes at log_2_(FC) >|2| and adjusted *p* <0.01) but not by Δ*dhbA* (9 differentially regulated genes). Approximately 4−8% of the transcriptomes of DK1042 and Δ*dhbA* (364 and 172 differentially regulated genes, respectively) were differentially regulated by the presence of PS92, but comparison of cocultures revealed that 1002 transcripts were differentially regulated between the two strains in coculture. In monoculture, 395 transcripts were differentially regulated between DK1042 and Δ*dhbA*.

### Δ*dhbA* downregulates sporulation and biofilm formation

When coculturing DK1042 with PS92, a large number of genes related to sporulation and germination were upregulated in DK1042 (Fig. 4a), though not differentially regulated in the Δ*dhbA* mutant in coculture compared with its monoculture. However, comparing the levels of transcription between DK1042 and Δ*dhbA* reveals that lack of *dhbA* strongly downregulated sporulation and germination pathways, independently of whether it was cocultured with PS92.

**Fig 4.**
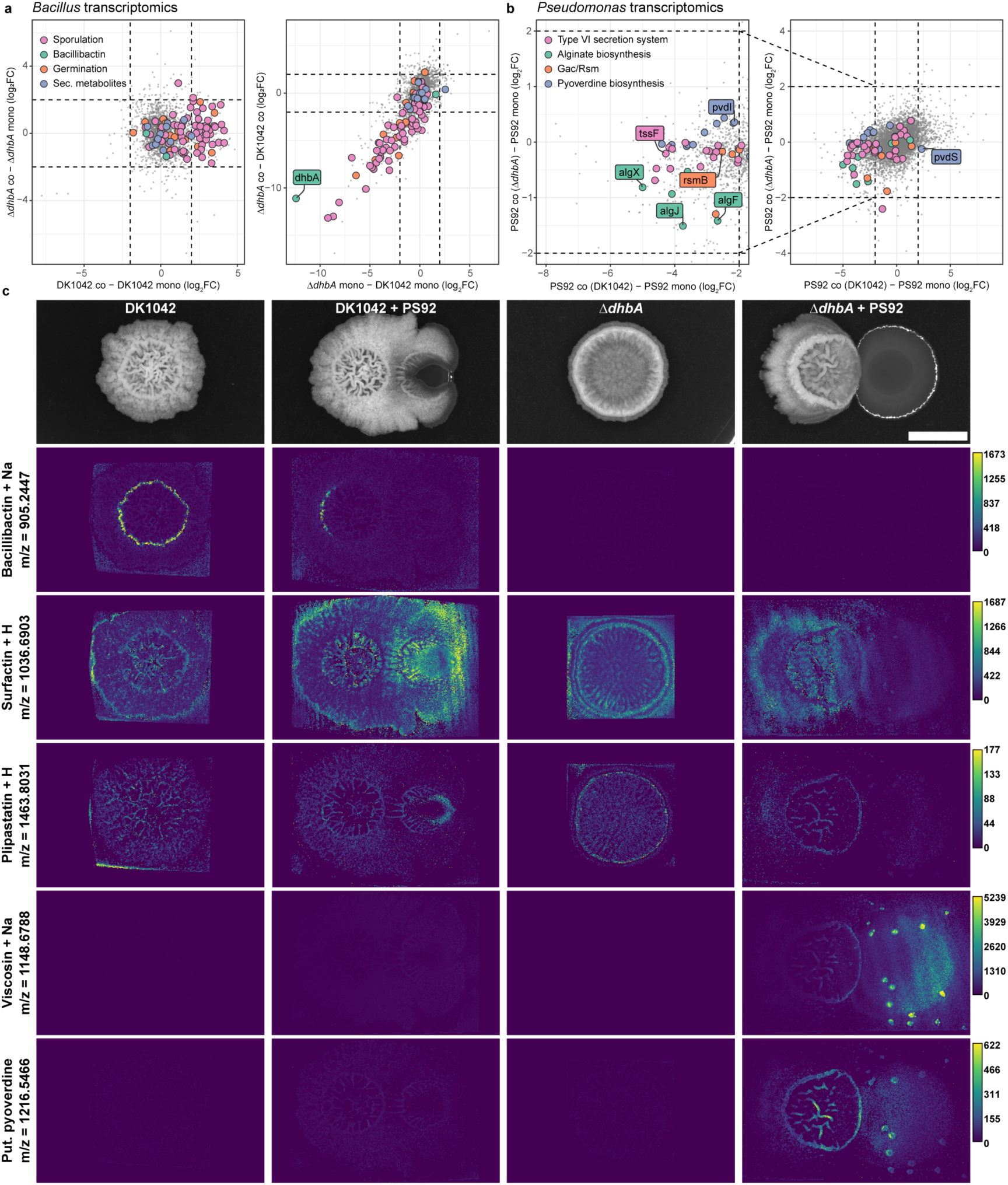
DK1042 alters PS92 transcriptome and metabolome via *gacS*. **a:** Δ*dhbA* downregulates genes related to sporulation and germination independently of having PS92 as its interaction partner. DK1042-upregulated genes of the same category only when in coculture with PS92. Dashed lines indicate log2(FC) = |2|. **b:** Several biosynthetic pathways related to the Gac/Rsm two-component system were significantly downregulated (log_2_(FC) <-2 and adjusted *p*-value <0.01) in coculture with DK1042, except for the alternative sigma factor *pvdS*. **c:** Mass spectrometry imaging reveals the presence/absence of select metabolites as well as their spatial localisation in the interactions. Scale bar = 5 mm. Grey values are root mean squared intensity across all samples, and lookup tables were scaled identically across each metabolite. NaN pixels were set to zero. Metabolites were annotated with metaspace using the 2019 Natural Product Atlas database. m/z = 1216.5466 was manually annotated as a pyoverdine structure.

Elimination of DHB and BB production affected several genes related to biofilm formation and sporulation in the *epsA-O*, *tapA-tasA-sipW*, *spsA-L*, *cot* and *spoII*, *III*, *IV*, *V* and *VI* operons, an effect reported previously [9, 51]. In addition, the transcription level of the *sinI* gene was higher in monoculture for Δ*dhbA* compared with DK1042, though not in coculture. In coculture, lack of *dhbA* upregulated the transcription of the biofilm repressor gene *abrB*. Together, this suggests that the Δ*dhbA* mutant did not form a biofilm or undergo sporulation either in coculture or in monoculture.

Repression of sporulation and biofilm formation was further corroborated by the downregulation of genes involved in utilisation of glutamine, manganese, and glycerol. Manganese and glutamine/glutamate utilisation are well known for their role in the Spo0A sporulation pathway that drives both biofilm formation and sporulation [52–55], while manganese homeostasis was very recently shown to be tightly interlinked with iron homeostasis [56].

Interestingly, the transcriptome of the Δ*dhbA* mutant did not display signs of iron starvation, with the exception of a slight repression in the transcription of *fur*. The *dhb*, *feu*, *yusV*, *besA,* and *yvm* genes were not differentially regulated in the Δ*dhbA* mutant compared with DK1042, except the *dhbA* transcript level which was deleted in the genome of the mutant and therefore transcriptionally absent in the mutant. Additionally, genes involved in production of known bioactive secondary metabolites were not differentially regulated in *Bacillus*, regardless of mutation or coculture partner.

### PS92 downregulates Gac/Rsm and iron acquisition in coculture with DK1042

When cocultured with DK1042, PS92 underwent downregulation of biosynthetic genes responsible for production of several secondary metabolites (Fig. 4b and Dataset S1). All biosynthetic genes involved in Pvd, alginate and viscosin production were repressed along with a Type VI secretion system (T6SS) subtype i3, while genes responsible for type II and III secretion and reactive oxygen neutralisation were upregulated. Similar changes have previously been observed in the closely related *P. fluorescence* SBW25 when removing the histidine kinase *gacS* (OrthoANIu similarity between SBW25 and PS92 = 99.06%) [57]. The Gac/Rsm pathway is a conserved two-component system found in many γ-proteobacteria and known to positively regulate biofilm formation and secondary metabolism in most pseudomonads and induce virulence factors in plant pathogenic species. Indeed, PS92 *gacS* (FNPKGJ_17835) was downregulated 4-fold in PS92 when cocultured with DK1042 but not when cocultured with Δ*dhbA*.

Unexpectedly, while Pvd biosynthetic genes were downregulated, the iron starvation sigma factor-encoding gene *pvdS* (FPNKGJ_22055) had a higher transcript level in coculture with DK1042. Pvd is the primary siderophore in SBW25 [58], though other iron-chelating molecules are encoded in the genomes of both SBW25 [57] and PS92. A biosynthetic gene cluster encoding the production machinery for a corrugatin-like molecule (FNPKGJ_15100-15175) was similarly repressed, although not below the threshold of log_2_(FC) <-2. Upregulation of *pvdS* suggests that PS92 is starved for iron in coculture with DK1042, but with downregulation of the iron acquisition systems it is unclear whether the consequence of coculture is iron starvation, repression of the Gac/Rsm pathway, or a combination of both.

We found further evidence for repression of the Gac/Rsm system in mass spectrometry imaging of cocultures showing that several molecules produced by genes under control of the Gac/Rsm system were absent from cocultures with DK1042 but present in monoculture colonies and cocultures with Δ*dhbA* (Fig. 4c). The presence of viscosin in all cultures except coculture with DK1042 on KB prompted us to test if viscosin was the antagonistic molecule against Δ*dhbA.* In lieu of a PS92 Δ*viscA* gene deletion mutant, we obtained a *P. fluorescens* SBW25 Δ*viscA* mutant with an identical interaction with DK1042, but found that *ΔviscA* was still able to antagonise Δ*dhbA*, similarly to the WT strain (Fig. S6).

### Antagonism depends on bacillibactin

Based on the transcriptome data for DK1042, we screened multiple gene disruption mutants for their antagonistic effect against PS92 (Fig. 5). Effects of mutations ranged from deficiency in production of biofilm matrix (*epsA-O*, *tasA*) and secondary metabolites (*srfAA*, *pksL*, *sfp*, *cypX*, *yvmC*) to lacking sensor kinases (*kinA-E*) or being unable to undergo sporulation (*spo0A*). Interestingly, only three mutants were significantly unable to antagonise PS92 (Δ*sfp*, Δ*feuA* and Δ*glpK*). Thus, neither sporulation nor biofilm formation are essential for antagonism, and likewise, typical secondary metabolites with antimicrobial properties such as surfactin and bacillaene had no effect on the restriction of PS92 colony area (although this was suggested from the mass spectrometry imaging). The inability of Δ*feuA* to antagonise suggests that iron uptake by BB is essential for restriction of PS92 colony size, but the fitness of the Δ*feuA* mutant was rather low even in monoculture (**Fig. S7**), which could result in low metabolic output in general. The antagonistic activity of the Δ*besA* mutant suggests that Fe(III) from BB is not required for antagonism, and that BB itself may inhibit PS92.

**Fig 5.**
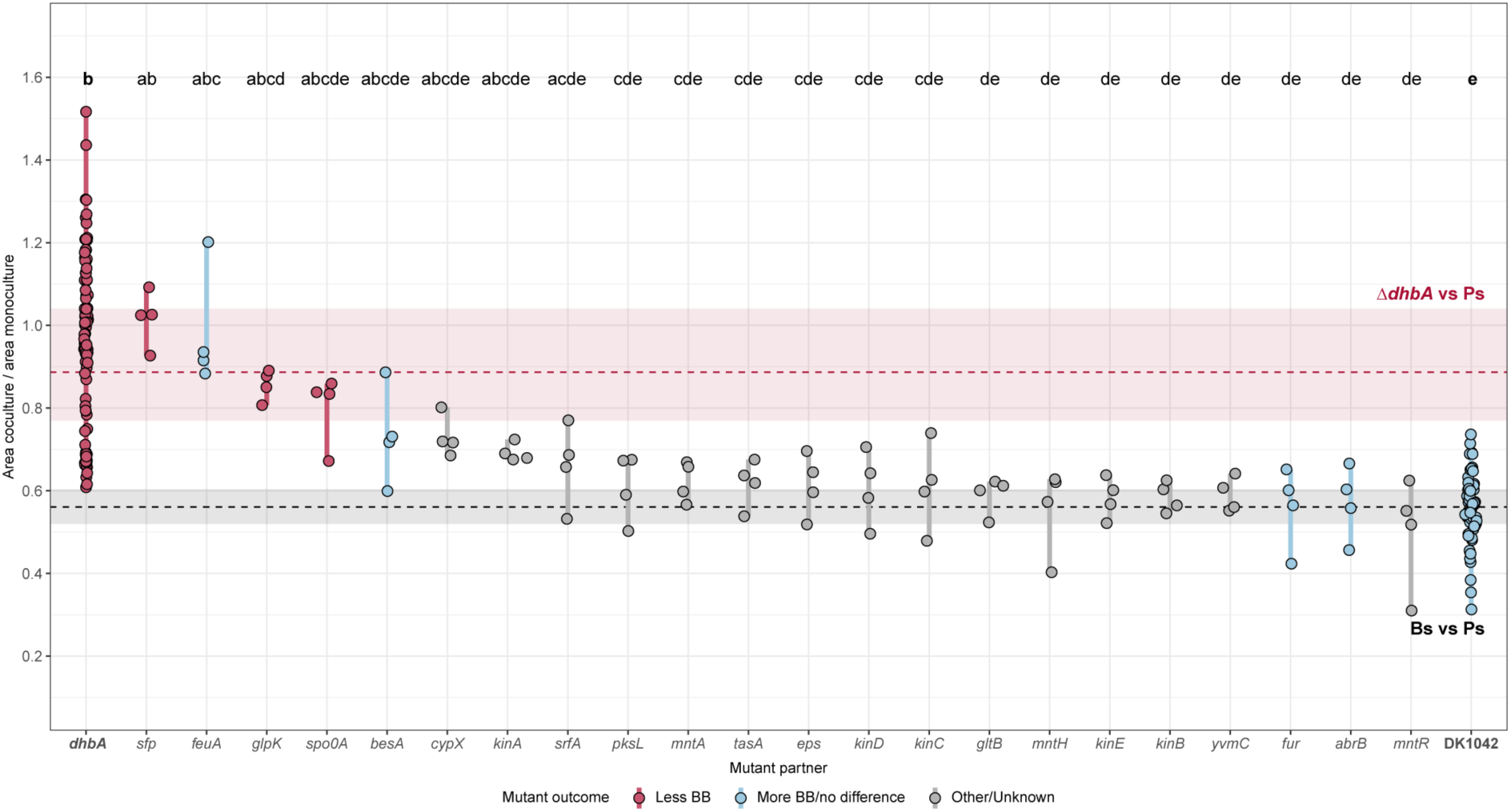
*Pseudomonas* colony size restriction is related to *sfp* and *feuA*. PS92 colony size was measured in cocultures next to *Bacillus* gene deletion mutants deficient in multiple aspects of cell physiology. Colonies spotted next to mutants lacking *sfp, feuA* and *glpK* were significantly larger than those spotted next to DK1042 relative to monoculture colony size, while not being significantly smaller than colonies spotted next to Δ*dhbA*. Other mutants related to biofilm formation, metal homeostasis and bioactive secondary metabolites were not diminished in their restriction of PS92. Dashed lines and rectangles are median, 1^st^ and 3^rd^ quartiles for PS92 colonies spotted next to Δ*dhbA* (red) and DK1042 (grey). Grouping letters are from ANOVA with Tukey-Kramer’s Post-hoc test. Identical letters within each plot indicate a statistically significant grouping (*p* <0.05, n = 4 biologically independent colonies). *dhbA* and DK1042 represent pooled data from multiple experiments (n >80).

### Interaction is conserved across fluorescent *Pseudomonas* isolates

To determine if the iron-related antagonism is specific to DK1042 and PS92, we cocultured DK1042 and Δ*dhbA* on KB agar with a collection of 16 fluorescent *Pseudomonas* soil isolates for which genomes had been sequenced (Fig. 6). Thus, we could qualitatively compare the interaction outcome with the genomic presence of biosynthetic pathways. Multiple isolates were restricted in colony size by DK1042 and not by Δ*dhbA*. Most isolates also seemed to influence the size and/or morphology of Δ*dhbA*, except XL272 (previously *Pseudomonas stutzeri*, now *Stutzerimonas degradans*) which is the only isolate lacking the Gac/Rsm-controlled biosynthetic machineries producing pyoverdine and alginate. A few isolates belonging to *Pseudomonas corrugata* and *protegens* subgroups were able to inhibit both DK1042 and Δ*dhbA*, which may be attributed to their potential for producing 2,4-diacetylphloroglucinol and pyoluteorin [59]. Strains from other *Pseudomonas fluorescens* subgroups were all restricted in colony growth in the presence of BB-producing *B. subtilis*, demonstrating the conservation of this antagonism across species.

**FIG 6.**
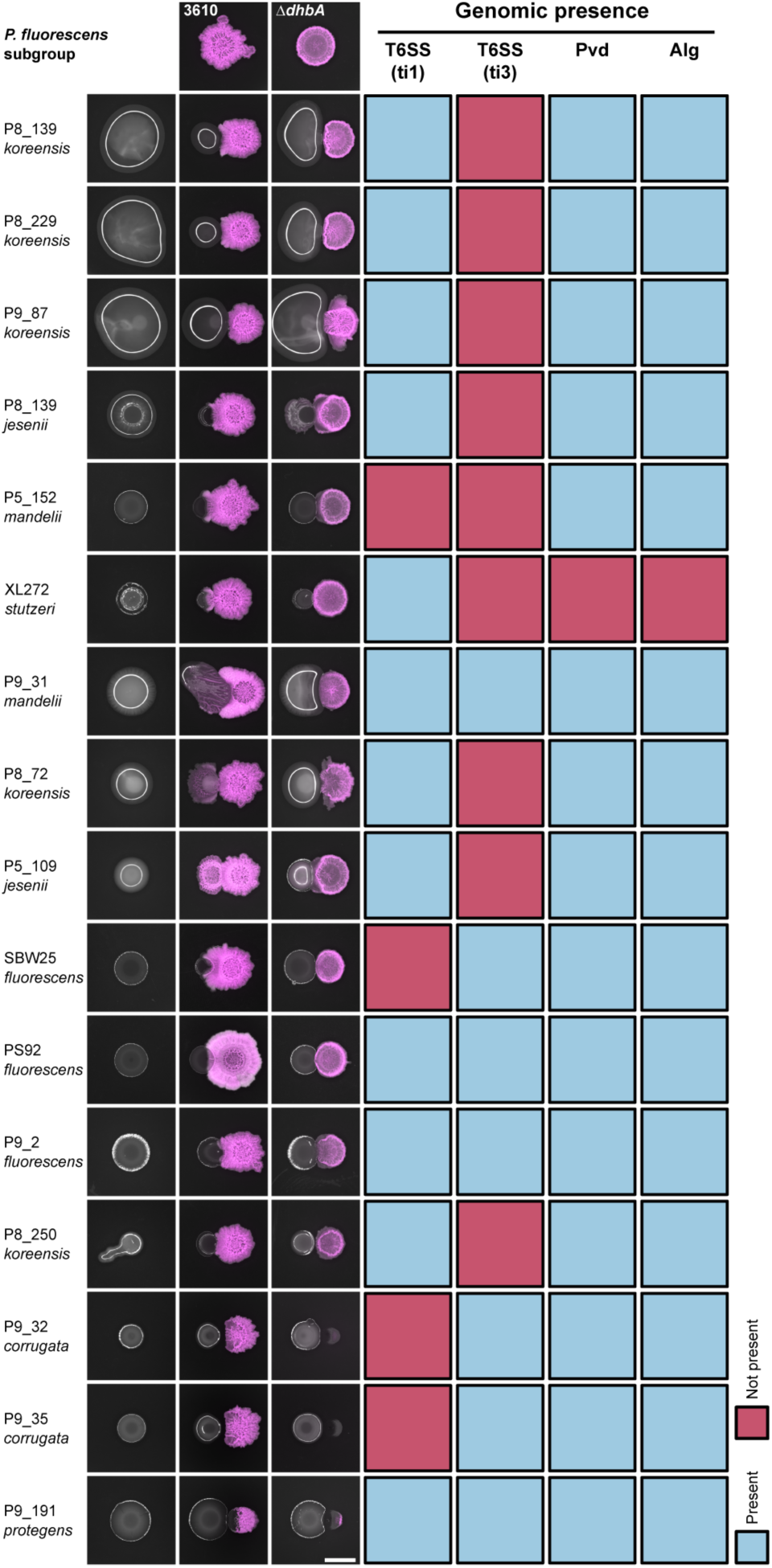
Antagonism is conserved across *Pseudomonas* spp. Fluorescent *Pseudomonas* soil isolates were cultured next to DK1042 or Δ*dhbA* on KB agar and their genomes were assessed for the presence of Gac/Rsm-related biosynthetic genes. Presence of >80% of PS92 genes at 40% amino acid identity was scored as ‘present’ (blue) while any less was scored as ‘not present’ (red). Isolates are presented as their strain ID and the closest related *P. fluorescens* subgroup as described by Hesse et al. (2016) based on whole-genome alignments. T6SS, Type VI Secretion system; ti1, Type i1; ti3, Type i3; Pvd, Pyoverdine; Alg, Alginate.

## DISCUSSION

In this study, we report a novel molecular interaction conserved between *B. subtilis* and multiple *Pseudomonas* spp. We show that the interaction is dependent on iron and BB, and that a lack of BB on KB agar results in both diminished offense and defence. The consequence of the BB-mediated antagonism is a restriction of *Pseudomonas* colony spreading and a reduction both in transcription and production of several products related to the Gac/Rsm system, which over time allows *Bacillus* to overgrow a *Pseudomonas* colony.

*Bacillus* defence is especially affected by iron availability. Iron homeostasis, biofilm formation, and sporulation are tightly interlinked in *B. subtilis*, and DhbA is essential for structured biofilm formation and proper sporulation due to the high iron demands of these lifestyles [9, 51, 60]. Surprisingly, matrix and sporulation as defensive measures in microbial ecology have received little attention [61], though the relation to T6SS has been investigated [24, 62]. Defence appears to take precedence for *Bacillus* in the interaction, and a biofilm- and sporulation-deficient mutant may be too antagonised by PS92 to produce the necessary effector for growth restriction. This could explain our observations with Δ*feuA* which requires large quantities of supplemented iron for proper fitness. Additionally, a biofilm-proficient strain lacking the antagonistic molecule may block PS92 growth through spatial competition [63–65], though it will be unable to restrict colony spreading chemically.

Another possibility could be that BB is a stronger iron chelator than Pvd, sequestering Fe(III) away from *Pseudomonas* and thereby inducing iron starvation. When *Bacillus* is unable to produce BB the tide turns, and Pvd from *Pseudomonas* sequesters Fe(III) away from *Bacillus*. Pvd production on KB has been shown in several pseudomonads to be highly dependent on iron-content, where iron-binding Fur represses Pvd production through repression of *pvdS* [58]. This could explain how Δ*dhbA*, Δ*besA* and Δ*feuA* are rescued by iron supplementation even in the form of ferric chloride. With this mechanism, strong iron starvation responses are likely in both *Bacillus* and *Pseudomonas*, but in Δ*dhbA* iron homeostasis was not significantly upregulated in coculture with PS92. On the other hand, *pvdS* in PS92 was significantly upregulated, which is indicative of iron starvation, but without the complementary upregulation of the rest of the pyoverdine biosynthetic genes. In *P. aeruginosa* Gac/Rsm is known to positively regulate pyoverdine biosynthetic gene transcription trough PvdS [66, 67], and a similar phenotype has been described in *P. fluorescens* SBW25 [57]. Recently, both pyoverdine and, to a larger extent, pyochelin were shown to be sensed by *B. velezensis*, instigating mobilisation of secondary metabolites [21]. Although sensing of pyochelin occurs independently of iron and BB, pyoverdine was shown to increase the production of BB. PS92 does not contain a gene cluster encoding the synthetic machinery for any type of pyochelin, and thus, pyochelin sensing presumably does not occur in the interaction between DK1042 and PS92.

Our results suggest *Bacillus*-mediated repression of *gacS*, a phenomenon with precedent from interaction between a *Bacillus* sp. isolate and *Pseudomonas syringae* pv. tomato DC3000 [68], though neither previous nor our experiments reveal if *gacS* repression is directly caused by DK1042 or by a sensory mechanism in PS92. Currently, there is no consensus on exactly which ligand triggers the Gac signal cascade [69], though it is expected that secreted products resulting from the activated pathway itself are at least partly responsible [70]. However, in *Pseudomonas* GacS is activated and inhibited by LadS and RetS, respectively, which in turn are activated by calcium ions (LadS) and mucin glycans or temperature (RetS) [71–73]. It is also hypothesised that the Gac/Rsm pathway functions via an alternative quorum sensing pathway, and that induction is achieved at higher cell densities [70]. This might explain why *gacS* was downregulated in PS92 spotted next to DK1042, where the colony is indeed restricted in size and at a lower cell density. Unfortunately, large-scale purification and chemical synthesis of BB was unsuccessful, hence we could not probe the antagonistic properties of BB directly.

Siderophores have long been implicated in microbial interactions, particularly in iron-scarce environments. In infections, pyoverdine and enterobactin are essential for *P. aeruginosa* and *E. coli*, respectively, not only due to iron requirements in a complex environment, but also in polymicrobial interactions as interspecies competitive factors [74]. Siderophores are also thought to play a role in biocontrol of plant pathogens by both *Pseudomonas* and *Bacillus*, but a thorough examination of the literature reveals that *Bacillus* and *Pseudomonas* siderophores have only rarely been directly implicated in interspecies interactions *in planta* [13]. Most evidence originated in the previous millennium where *Pseudomonas* siderophores proved to be determinants of microbial interactions through iron sequestration [75, 76]. Others have since inferred biocontrol potential from the ability to produce siderophores without experimentally determining the consequence of losing siderophore production [10, 77]. Our study systematically investigated the significance of BB and its effects on transcription and metabolite production in microbial interactions. Although we demonstrated the importance of BB on rich media, our study complements and expands the previously established role of BB in biocontrol. Our *P. marginalis* isolate proved non-pathogenic against lettuce, cabbage and spinach, hence we were unable to determine if BB-mediated antagonism occurs in the phyllosphere as well as on agar surfaces. Other studies have successfully transferred biocontrol properties from *in vitro* to *in planta* [78], but the medium dependency of the antagonism between DK1042 and PS92 might render it difficult to apply to agriculture. Regardless, understanding the molecular mechanism underlying the antagonism could allow the environment to be tailored specifically to the intended interaction. Future studies should investigate the apparent tug-of-war between BB and Pvd, as well as the hypothesis that BB represses Gac sensing in fluorescent pseudomonads.

## ACKNOWLEDGEMENTS

The authors thank Lars Jelsbak and Morten Lindquist Hansen for contributing *Pseudomonas* soil isolates for this study, and the DTU Metabolomics Core for MSI and LC-MS instrumentation. This project was funded by a DTU Alliance Strategic Partnership PhD fellowship, by the Danish National Research Foundation (DNRF137) for the Center for Microbial Secondary Metabolites, and the Novo Nordisk Foundation within the INTERACT project of the Collaborative Crop Resiliency Program (NNF19SA0059360) and for the “Imaging microbial language in biocontrol (IMLiB)” infrastructure grant (NNF19OC0055625).

## AUTHOR CONTRIBUTIONS

ML and ÁTK designed the study; ML, JPBJ, and SAJ collected and analyzed data; MDS, DKCA, and CNLA contributed with methodology or preliminary data; ML and ÁTK wrote first draft of paper with edits from all authors.

## COMPETING INTERESTS

The authors declare no competing interests.

## Notes

### Competing Interest Statement

The authors have declared no competing interest.

