## Supplementary materials for "Competition for iron shapes metabolic antagonism between *Bacillus subtilis* and *Pseudomonas*"

**Video S1: Interaction timelapse.** DK1042 (left, magenta) or Δ*dhbA* (right, magenta) were spotted next to PS92 (green) on King's B agar and incubated at 30 °C in a Zeiss Axio Zoom V16 stereomicroscope. Images were acquired every 15 minutes for 72h.

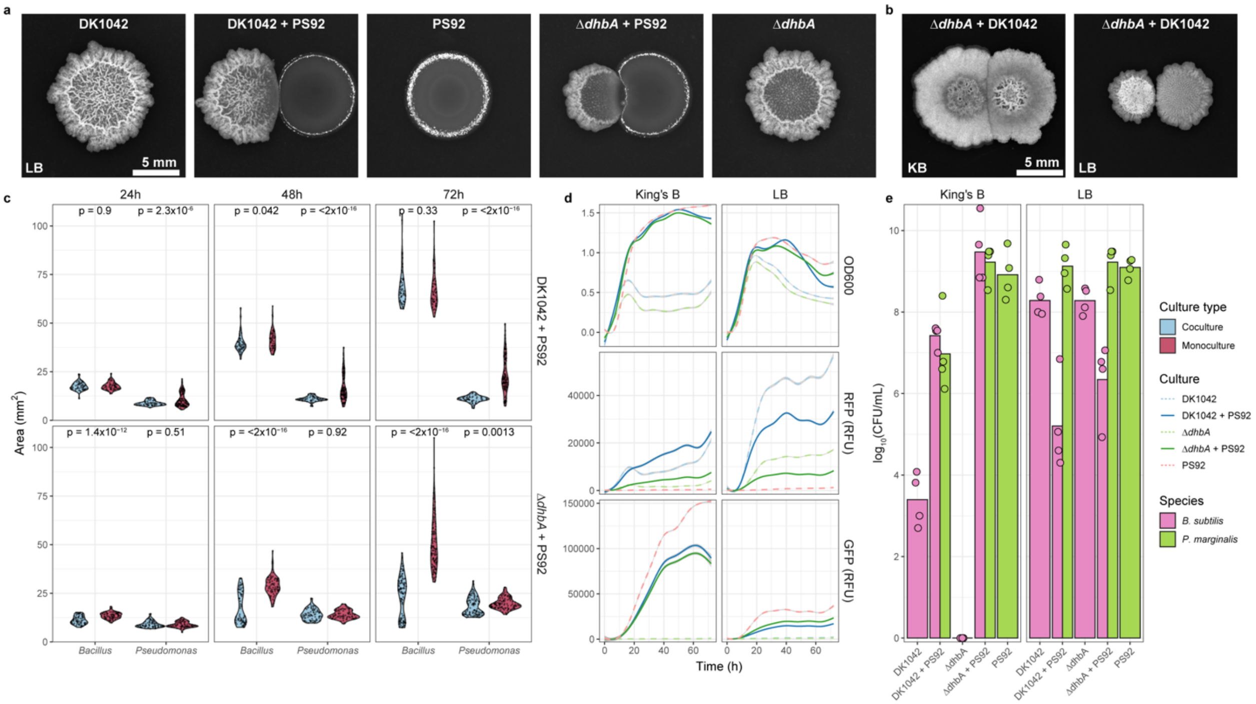


**Figure S1: Interaction is constrained to solid King's B. a:** Colonies grown on LB for 72h at 30 °C. **b:** Δ*dhbA* spotted next to DK1042 on King's B and LB and grown at 30 °C for 72h. **c:** Non-normalized colony-areas visualized in Figure 1. p-values are from Student's t-test adjusted for multiple testing by the Benjamini-Hochberg method. **d:** Growth in liquid King's B and LB. RFP: mKate2 signal (*Bacillus*), GFP: msfGFP signal (*Pseudomonas*). **e:** Viable cell count from planktonic cultures.

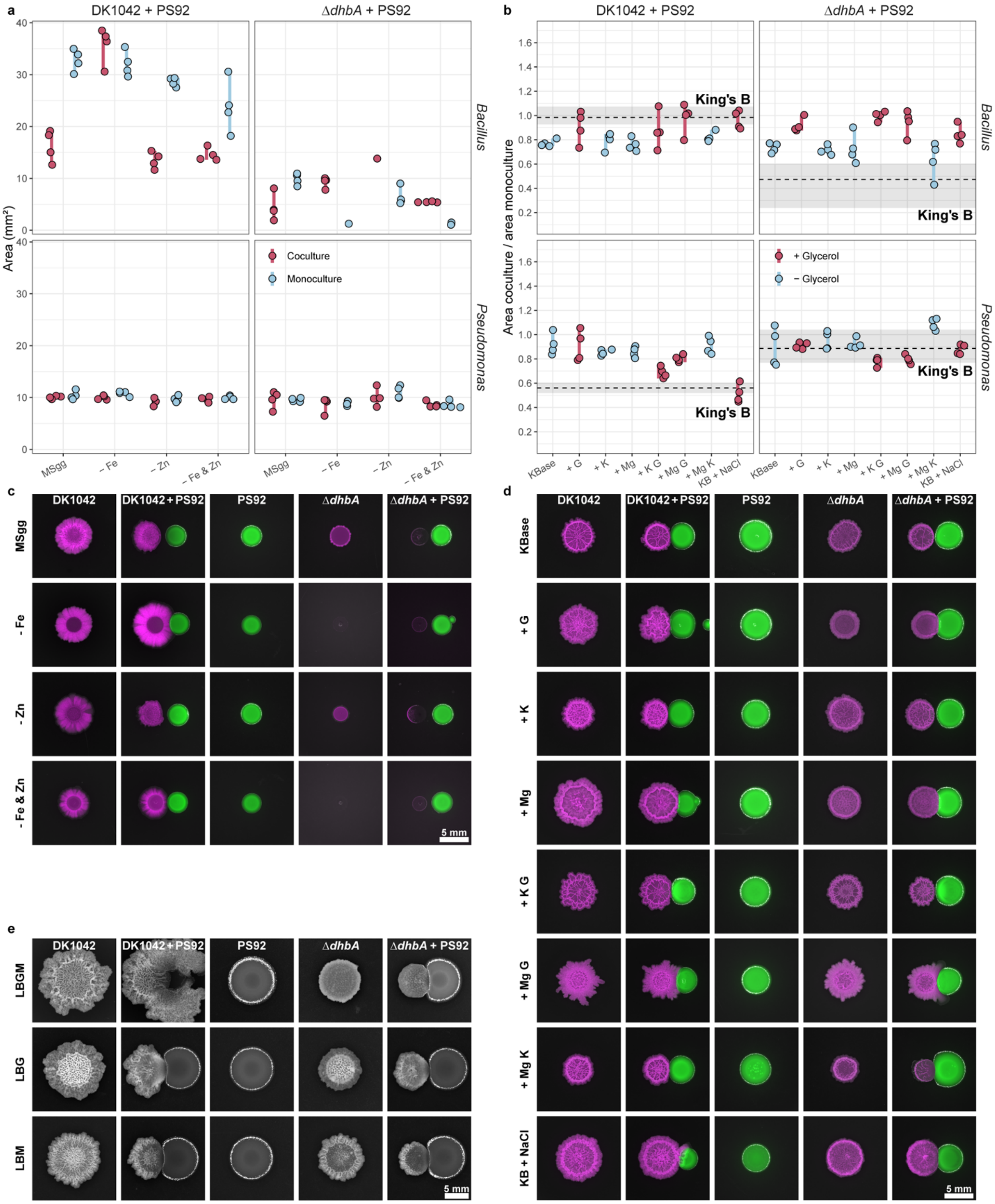


**Figure S2: Interaction depends on medium composition. a:** Colony area of *Bacillus* and *Pseudomonas* in interactions on MSgg with and without supplemented metals. Interactions were cultivated at 30 °C for 72h. **b:** Relative colony size of *Bacillus* and PS92 cultivated on King's B with or without supplemented $Mg^{2+}$, $K^{+}$, or glycerol (G). KBase is water and 20 g/L peptone. Dashed lines and squares are median, 1st, and 3rd quantile, respectively for DK1042 + PS92 (left) and Δ*dhbA* + PS92 (right) on regular King's B. **c and d:** Stereomicroscopy of representative samples from **a** and **b.**

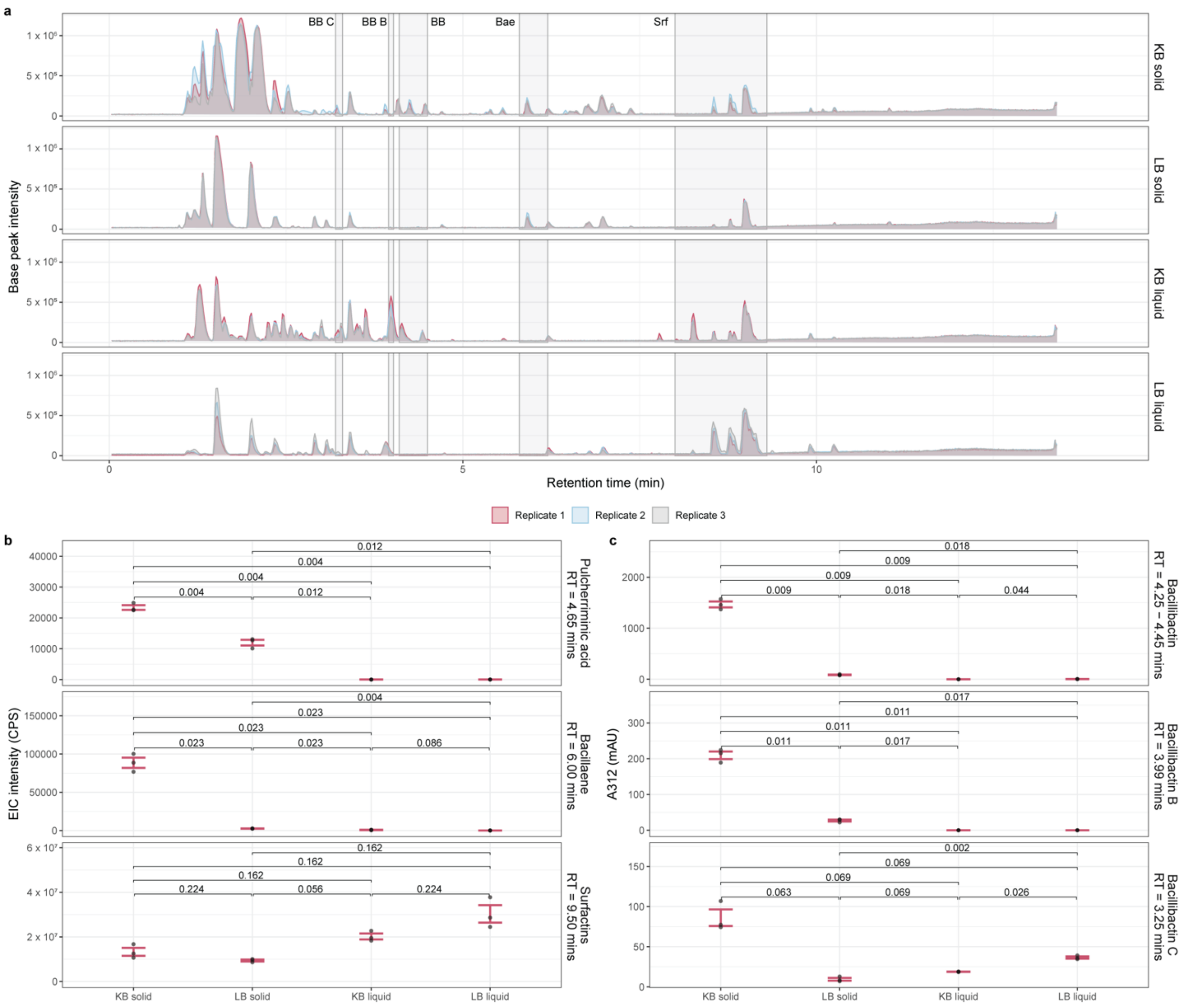


**Figure S3: LC-MS in solid and liquid King's B and LB. a:** Base peak chromatogram of DK1042 grown at 30 °C for 72h. **b:** Integrated peak area of select secondary metabolites measured in their respective ion chromatograms extracted by m/z-value. p-values are from Student's t-test adjusted for multiple testing by the Benjamini-Hochberg method. **c:** Same as **b**, but with Bacillibactins measured from their UV chromatograms.

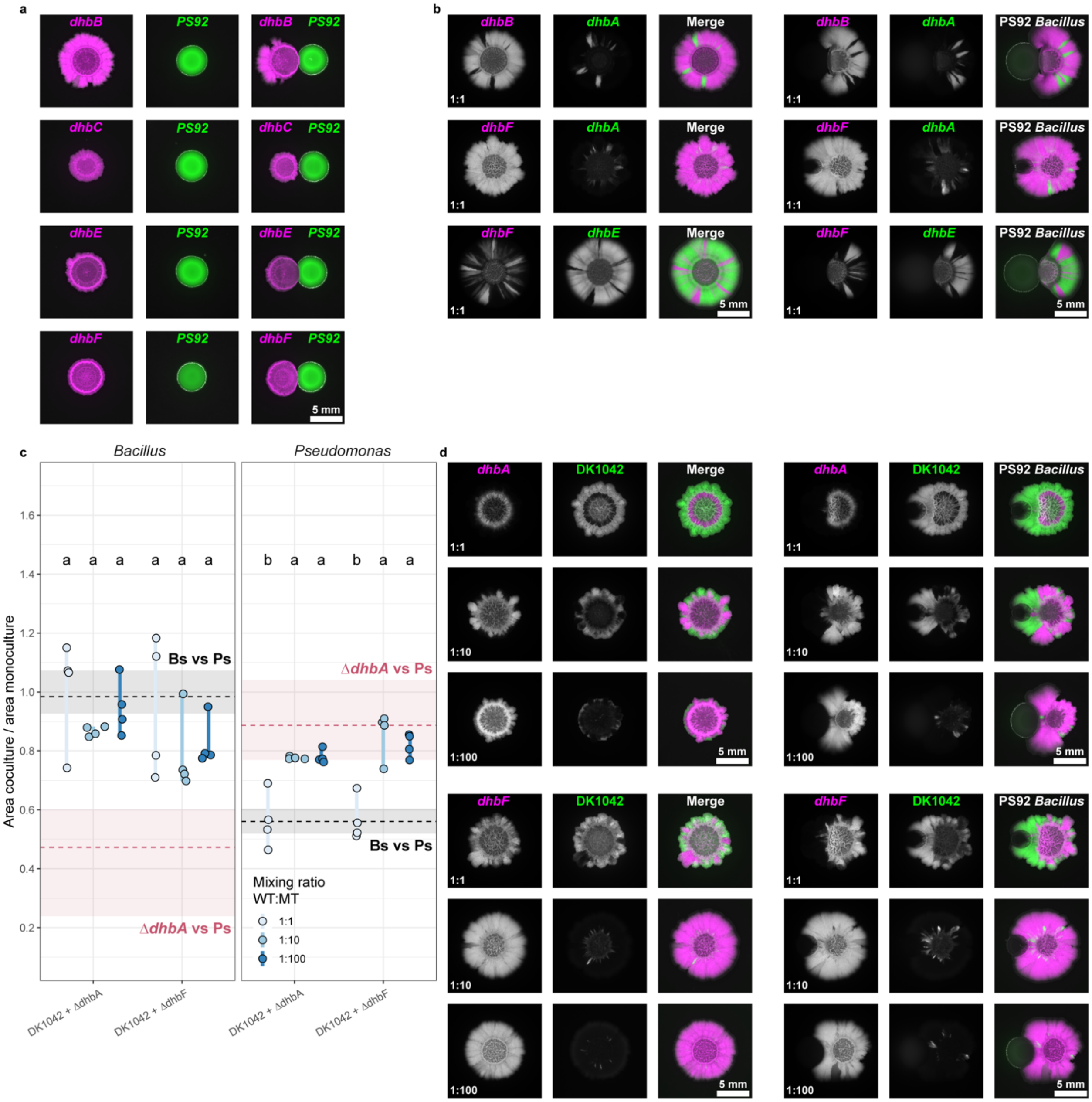


**Figure S4: DK1042 can complement bacillibactin-deficient mutants. a:** Stereomicroscopy of single gene deletion mutants deficient in genes encoded in the *dhb* operon cultivated on King's B at 30 °C for 72h either alone or next to PS92. See Figure 2b. **b:** Stereomicroscopy of *dhb* mutants mixed in equal ratios and their interactions as in **a**. See Figure 2c. **c:** Relative colony size of *Bacillus* and PS92 cultivated as **a** and **b**. Dashed lines and squares are median, 1st, and 3rd quantile, respectively for DK1042 + PS92 (grey) and Δ*dhbA* + PS92 (red). **d:** Stereomicroscopy of representative samples from **c**.

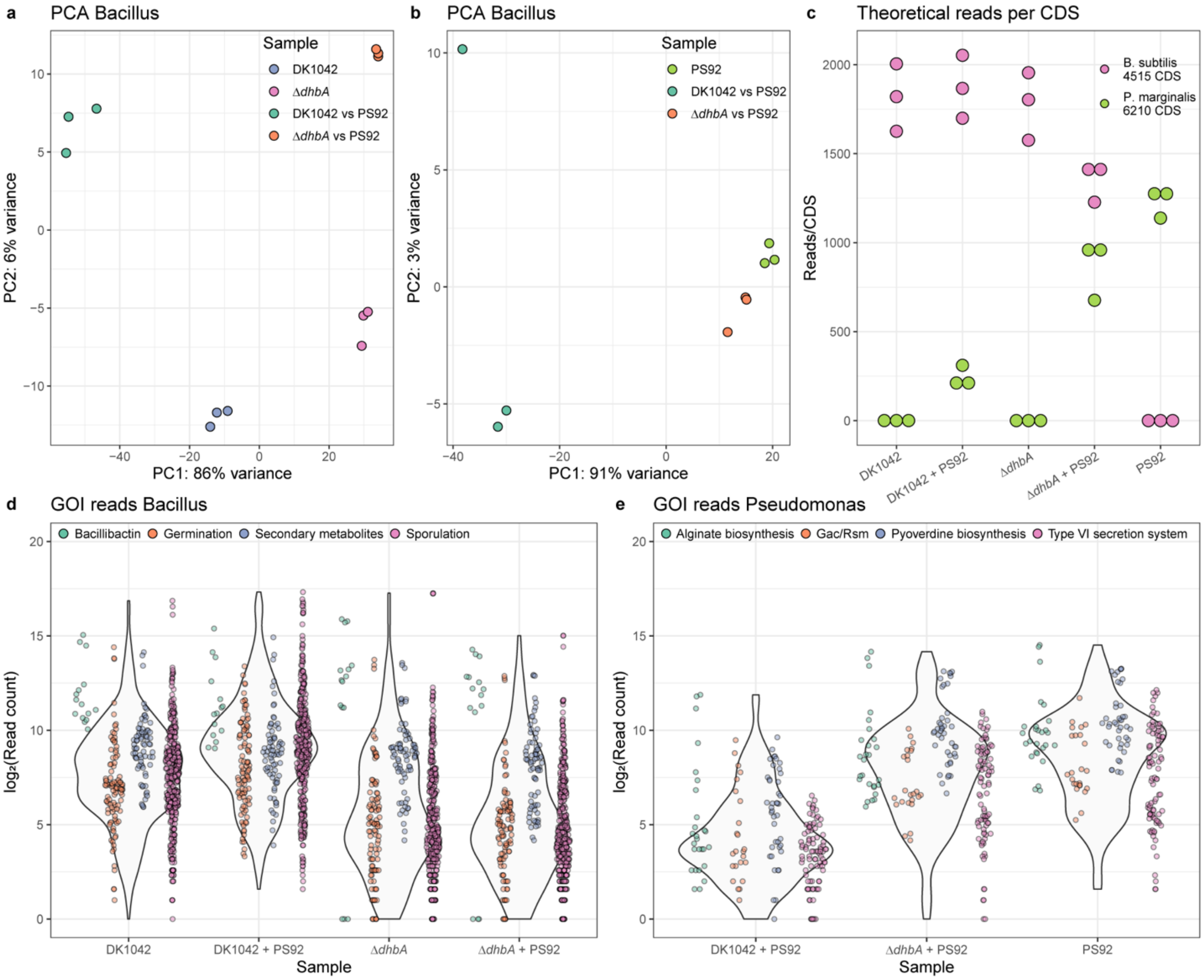


**Figure S5: RNAseq quality checks. a and b:** Principal component analysis of transcript reads in each sample for *Bacillus* (**a**) and *Pseudomonas* (**b**). **c:** Theoretical number of reads per coding sequence (CDS) in each genome for each sample. Calculated as $n_{reads}/n_{CDS}$. **d and e:** log-transformed read count for genes of interest highlighted in Figure 4a and 4b for *Bacillus* (**d**) and *Pseudomonas* (**e**).

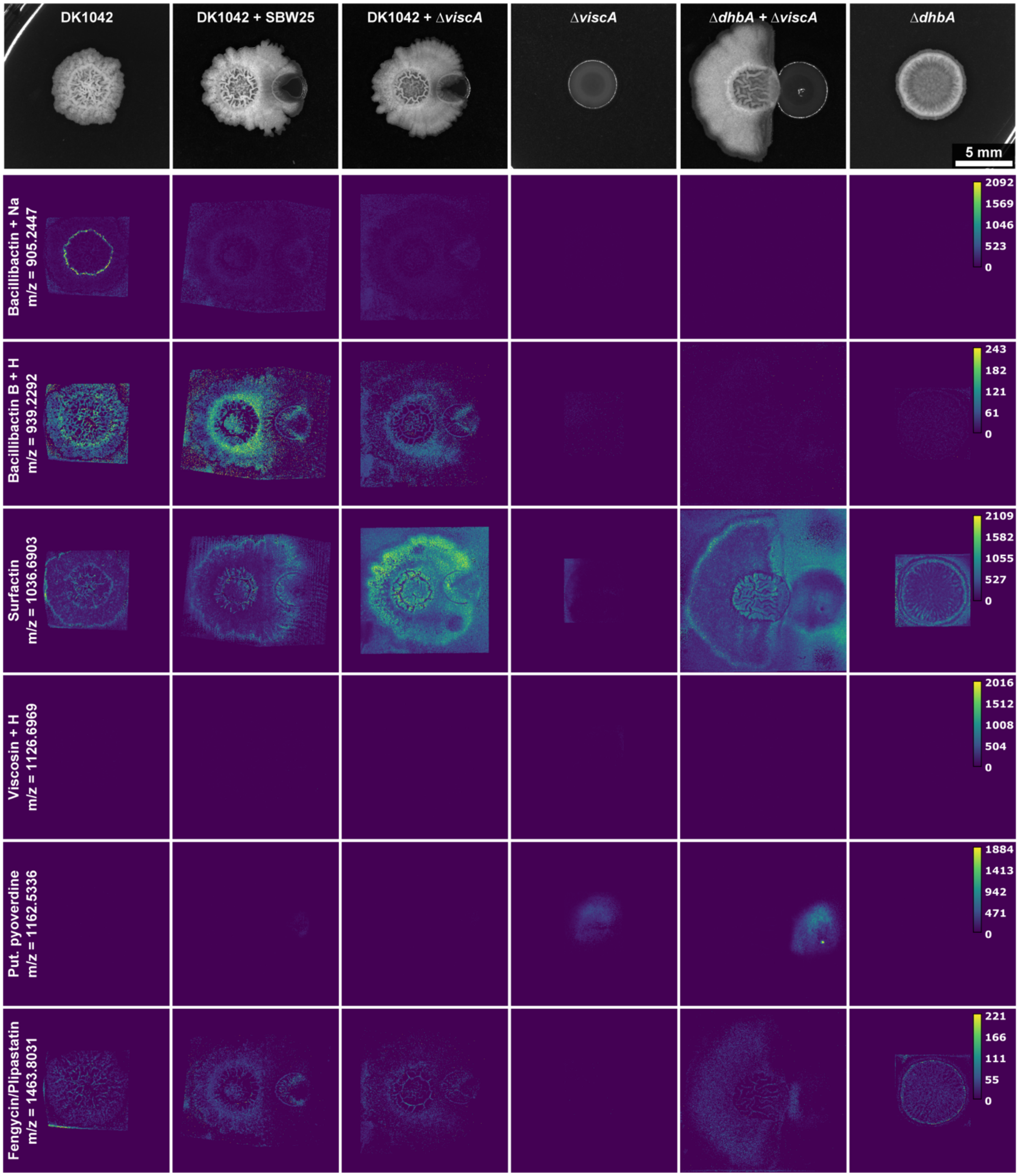


**Figure S6: Viscosin is not responsible for *Bacillus* antagonism.** Close-proximity colonies of DK1042 and *Pseudomonas fluorescens* SBW25 wild type and Δ*viscA* deficient in viscosin production. Colonies were cultivated on King's B at 30 °C for 72h. Mass spectrometry imaging reveals the presence/absence of select metabolites as well as their spatial localization in the interactions. Grey value is root mean squared intensity across all samples, and lookup tables were scaled identically across each metabolite. NaN pixels were set to zero. Metabolites were annotated with metaspace using the 2019 Natural Product Atlas database. m/z = 1162.5336 was manually annotated as a pyoverdine structure (this m/z-value is different from the pyoverdine structure in figure 4).

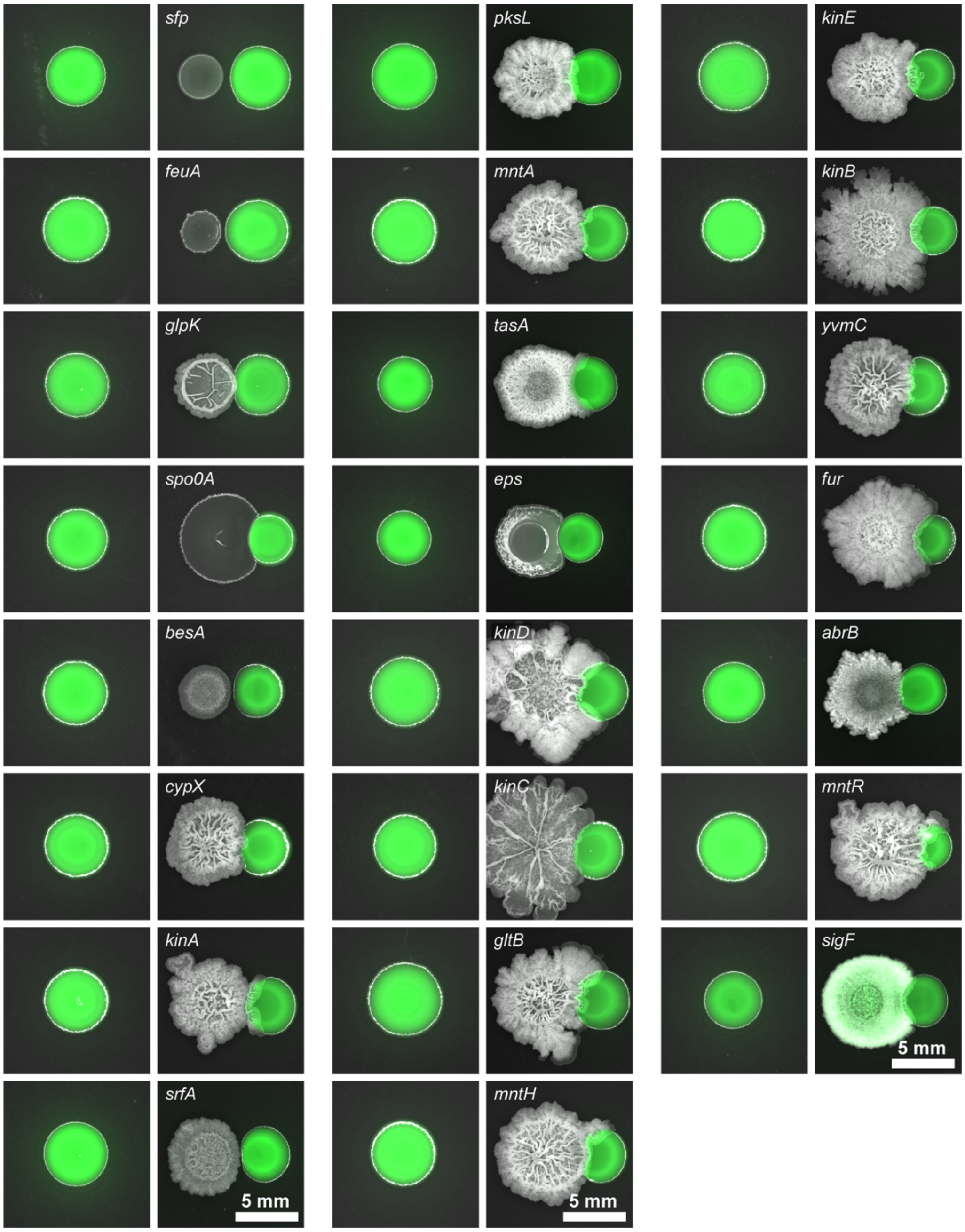


**Figure S7: *Bacillus* mutant interactions with PS92.** Stereomicroscopy of representative mutant interactions from Figure 5. Green is PS92.

**Table S1. Strains and oligos used in the study.**

| **Strain** | | | |
|---|---|---|---|
| ***Bacillus subtilis*** | **Genotype** | | **Ref** |
| *B. subtilis* DK1042 | 3610 *comI*$^{Q12I}$ | | [1] |
| TB501.1 | 3610 *comI*$^{Q12I}$ *amyE*::P$_{hyperspank}$-mKate2-*Spec*$^{R}$ | | [2] |
| TB500.1 | 3610 *comI*$^{Q12I}$ *amyE*::P$_{hyperspank}$-eGFP-*Spec*$^{R}$ | | [2] |
| Δ*dhbA* | DK1042 *dhbA*::*Kan*$^{R}$ | | This study |
| Δ*dhbA* mKate2 | DK1042 *dhbA*::*Kan*$^{R}$ *amyE*::P$_{hyperspank}$-mKate2-*Spec*$^{R}$ | | This study |
| Δ*dhbA* eGFP | DK1042 *dhbA*::*Kan*$^{R}$ *amyE*::P$_{hyperspank}$-eGFP-*Spec*$^{R}$ | | This study |
| Δ*dhbB* mKate2 | DK1042 *dhbB*::*Kan*$^{R}$ *amyE*::P$_{hyperspank}$-mKate2-*Spec*$^{R}$ | | This study |
| Δ*dhbC* mKate2 | DK1042 *dhbC*::*Kan*$^{R}$ *amyE*::P$_{hyperspank}$-mKate2-*Spec*$^{R}$ | | This study |
| Δ*dhbE* mKate2 | DK1042 *dhbE*::*Kan*$^{R}$ *amyE*::P$_{hyperspank}$-mKate2-*Spec*$^{R}$ | | This study |
| Δ*dhbE* eGFP | DK1042 *dhbE*::*Kan*$^{R}$ *amyE*::P$_{hyperspank}$-eGFP-*Spec*$^{R}$ | | This study |
| Δ*dhbF* mKate2 | DK1042 *dhbF*::*Kan*$^{R}$ *amyE*::P$_{hyperspank}$-mKate2-*Spec*$^{R}$ | | This study |
| DS3337 | 3610 Δ*sfp::MLS*$^{R}$ | | [3] |
| Δ*mntH* | DK1042 *mntH*::*Kan*$^{R}$ | | This study |
| Δ*mntA* | DK1042 *mntA*::*Kan*$^{R}$ | | This study |
| Δ*mntR* | DK1042 *mntR*::*Kan*$^{R}$ | | This study |
| DS4085 | 3610 *pksL*::*Cm*$^{R}$ | | [4] |
| DS1122 | 3610 Δ*srfA-C*::Tn10-*Spec*$^{R}$ | | [5] |
| Δ*gltB* | DK1042 *gltB*::*Kan*$^{R}$ | | This study |
| Δ*glpK* | DK1042 *glpK*::*Kan*$^{R}$ | | This study |
| TB863 | 3610 Δ*tasA::Km*$^{R}$ | | [2] |
| Δ*abrB* | DK1042 *abrB*::*Kan*$^{R}$ | | This study |
| Δ*cypX* | DK1042 *cypX*::*Kan*$^{R}$ | | This study |
| TB601 | 3610 Δ*epsA-O::Tet*$^{R}$ | | [2] |
| Δ*yvmC* | DK1042 *yvmC*::*Kan*$^{R}$ | | This study |
| TB398 | DK1042 *kinA*::*Erm*$^{R}$ | | This study |
| TB399 | DK1042 *kinB*::*Tet*$^{R}$ | | This study |
| TB400 | DK1042 *kinC*::*Spec*$^{R}$ | | This study |
| TB401 | DK1042 *kinD*::*Cm*$^{R}$ | | This study |
| TB402 | DK1042 *kinE*::*Cm*$^{R}$ | | This study |
| Δ*abrB* | DK1042 *abrB*::*kan*$^{R}$ | | This study |
| Δ*spo0A* | DK1042 *spo0A*::*Kan*$^{R}$ | | This study |
| Δ*fur* | DK1042 *fur*::*Kan*$^{R}$ | | This study |
| Δ*feuA* | DK1042 *feuA*::*Kan*$^{R}$ | | This study |
| Δ*feuA* mKate2 | DK1042 *feuA*::*Kan*$^{R}$ *amyE*::P$_{hyperspank}$-mKate2-*Spec*$^{R}$ | | This study |
| Δ*besA* | DK1042 *besA*::*Kan*$^{R}$ | | This study |
| Δ*besA* mKate2 | DK1042 *besA*::*Kan*$^{R}$ *amyE*::P$_{hyperspank}$-mKate2-*Spec*$^{R}$ | | This study |
| ***Pseudomonas*** | **Genotype** | **Genome Accession** | **Ref** |
| *P. marginalis* PS92 | WT isolate | CP125381 | This study |
| *P. marginalis* PS92 msfGFP | PS92 attTn7::P$_{14g}$-msfGFP-Gm$^{R}$ | | This study |
| *Pseudomonas* sp. P5_109 | WT isolate | CP125380 | This study |
| *Pseudomonas* sp. P5_152 | WT isolate | | This study |
| *P. zeae* P8_72 | WT isolate | | This study |
| *Pseudomonas* sp. P8_139 | WT isolate | CP125379 | This study |
| *Pseudomonas* sp. P8_229 | WT isolate | CP125378 | This study |
| *Pseudomonas* sp. P8_241 | WT isolate | CP125377 | This study |
| *Pseudomonas* sp. P8_250 | WT isolate | | This study |
| *Pseudomonas* sp. P9_31 | WT isolate | CP125375 | This study |
| *Pseudomonas* sp. P9_2 | WT isolate | CP125376 | This study |
| *Pseudomonas* sp. P9_32 | WT isolate | CP125374 | This study |

| | | | |
|---|---|---|---|
| *Pseudomonas* sp. P9_35 | WT isolate | CP125373 | This study |
| *P. germanicum* P9_87 | WT isolate | CP125372 | This study |
| *P. protegens* P9_191 | WT isolate | | This study |
| ***Escherichia coli*** | **Genotype** | | **Ref** |
| *E. coli* CC118 | CC118 λpir/pBG42 | | [6] |
| *E. coli* HB101 | HB101 /pRK600 | | [6] |
| *E. coli* CC118 | CC118 λpir/pTNS2 | | [6] |

| Oligo ID | Description | Sequence (5' – 3') |
|---|---|---|
| oTB134 | sfp up | GGTGTCAAGCTGTTGATGAG |
| oTB135 | sfp down | AAGCATCTCCGCCTGTACAC |
| oTB249 | feuA up | GCTGAGATGCTTTGCCCCAT |
| oTB250 | feuA down | CGAACGCATCGTTCAGCAAA |
| oTB251 | fur up | GCATCGACAGCTGTTTTCAG |
| oTB252 | fur down | GGAACTCTGCGCGTATTTTGT |
| oTB232 | dhbA up | CGATGAGGAGACGCTAAG |
| oTB233 | dhbA down | CGCTGGATGTCTGTTTGG |
| oTB255 | dhbB up | GTGCCGCAAAGCTTCTTCAA |
| oTB256 | dhbB down | CTGTCCATGATGCGGCAATG |
| oTB257 | dhbC up | AGCCCTTGTGTCAAGCTCTC |
| oTB258 | dhbC down | CCCGGATCAACGGAAACAGA |
| oTB259 | dhbE up | GATTCGGATCAGGCACCCAT |
| oTB260 | dhbE down | GGTTCAAAGCCTGAGGACGA |
| oTB261 | dhbF up | GTGGGCCGCTGTGTTTTTAG |
| | dhbF down | GCAGAGGTGACTTTCGTGGA |
| | abrB up | CGCAATGTACCAAAGCGTGA |
| | abrB down | GCTATGAAGGTAAGGATTTTGTCG |
| | glpK up | CTGCCAAGCTGGGTGTTTTC |
| | glpK down | CCGAATGTGCCGCCTTTATG |
| | gltB up | CGGCCTGTAATTGACCGGAT |
| | gltB down | CGCAAGCATCGAAGAGCAAA |
| | mntH up | AGCAGAACCGACCAAGAAGG |
| | mntH down | GACTTCTTGGCACATGGGGA |
| | mntA up | GGCTGATTTGCCTTTTCGCA |
| | mntA down | ACAGCCTGGTTTAGGGGTTG |
| | mntR up | GACCCTTTCTGCAAATCGCC |
| | mntR down | CGGATGCAGCGCAATTTCAA |
| | yvmC up | GATCCAGTGCGTCACCGATA |
| | yvmC down | TGACGTTTGGAAGGAAAAGGGA |

TB398-TB402 were created by transforming DK1042 with genomic DNA from JH12638 (*kinA*), JH19980 (*kinB*) from Wang *et al.* 1997 (Genes Dev), and BAL393 (*kinC*), BAL691 (*kinD*), and BAL692 (*kinE*) from Grau *et al.* 2015 (mBio).
